# Motor protein-assisted glycan translocation and sequencing using nanopores

**DOI:** 10.1101/2024.11.27.625763

**Authors:** Wenjun Ke, Pengcheng Wu, Shengzhou Ma, Yuting Yang, Yiwei Zhang, Qingqing Zhang, Xiaojie Lu, Bingqing Xia, Liuqing Wen, Jingwei Bai, Zhaobing Gao

**Affiliations:** State Key Laboratory of Drug Research, Shanghai Institute of Materia Medica, Chinese Academy of Sciences, Shanghai, 201203, China; University of Chinese Academy of Sciences, Beijing, 100049, China; School of Pharmaceutical Sciences, Tsinghua University, Beijing 100084, China; Tianjin University of Traditional Chinese Medicine, Tianjin 301617, China

## Abstract

The glycan sequencing remains a significant bottleneck in glycoscience. While nanopore platforms have achieved substantial progress in single-molecule nucleic acid sequencing, their application to glycan sequencing has faced considerable challenges, with limited advancements to date. In this study, we propose a novel strategy for controlling glycan translocation through the MspA nanopore as an initial step toward glycan sequencing. By conjugating the target glycan with a helicase-controlled single-stranded DNA, we successfully achieved sequencing reads of up to eleven glycans. For the first time, we isolated glycan-associated electrical signals, enabling the translocation, stretching, and controlled speed of neutral glycans through the nanopore. This method provides a platform for obtaining glycan read lengths and identifying different glycan modifications, demonstrating the capability to resolve monosaccharide composition and glycosidic linkages. To further improve resolution, we propose engineered M2-MspA to reduce the pore constriction size and enhance precision by minimizing the random thermal motion of the translocating glycan. These modifications are expected to increase the sequencing accuracy and reliability. This work represents the first proof-of-concept demonstration of glycan chain nanopore sequencing and lays a promising foundation for the development of single-molecule glycan fingerprinting and sequencing technologies. We anticipate that this approach will significantly advance the development and commercialization of nanopore-based glycan sequencing techniques.

## Introduction

Nanopores have emerged as a highly sensitive, single-molecule sensing technology^1–4^, demonstrating promising potential for glycan detection^5, 6^. Recent advancements in nanopore technology have enabled the discrimination of various glycans by monitoring the changes in ionic current as molecules pass through the pore, opening new pathways for glycan structure sequencing^7–14^. These achievements have been validated across several studies, marking significant progress in the fundamental sensing of glycans. However, despite these advancements, the direct sequencing of glycan chains via nanopores faces considerable technical challenges.

Unlike nucleic acids, glycan biosynthesis is a non-template-driven process influenced by multiple factors^15–18^, resulting in structurally complex and heterogeneous molecules^19, 20^. This structural diversity complicates glycan sequencing, as it is currently impossible to deduce glycan composition and linkages from genetic templates alone. Additionally, the intricate nature of glycans prevents the application of standard amplification techniques, underscoring the need for a method capable of single-molecule resolution in detecting monosaccharide composition, linkages, and modifications. Such capabilities would greatly advance the fields of glycoscience and glycomics. Although single-molecule nanopore sequencing of proteins has not yet reached full maturity, pioneering work in this domain offers valuable insights for glycan sequencing^21–25^.

In this context, motor protein-driven nanopore sequencing emerges as a promising strategy due to its ability to guide molecular substrates and facilitate sequence-specific reads^26–28^. Motor proteins, such as DNA helicases or polymerases, can bind target molecules and drive them progressively through the nanopore, generating distinct current signatures that reflect structural changes^29–31^. This approach has been widely applied in nucleic acid sequencing^32–34^ and has recently shown potential in protein sequencing, with the capability to detect single amino acids, modifications, and read lengths exceeding 100 kDa^22–25^. Nevertheless, the unique challenges of glycan sequencing exceed those encountered with nucleic acids and proteins. First, most glycans are electrically neutral, lacking the electrostatic driving force essential for pore translocation^35^. Additionally, the branched and sterically hindered structures of glycans impede their smooth passage through the nanopore. Achieving precise control over the translocation speed is another technical bottleneck crucial for high-resolution glycan sequencing. Finally, the structural heterogeneity of glycans can lead to irregular current signals, posing further challenges to data accuracy and completeness. These obstacles highlight why, to date, no successful reports exist for motor protein-driven nanopore glycan sequencing.

Here, we present the first proof-of-concept exploration of DNA motor proteins, such as helicases or polymerases, to investigate their potential in glycan sequencing. Specifically, we aim to demonstrate (1) the feasibility of glycan extension within the nanopore and the isolation of glycan-specific signals during translocation, establishing the viability of motor protein-driven nanopore glycan sequencing; (2) the correlation between glycan composition and electrical signatures; and (3) the capability of nanopore-based ionic current reads to reveal information on monosaccharide composition, linkages, and modifications within a single read. We anticipate that these foundational studies will significantly advance the development of a single-molecule glycan sequencing platform.

## Results

### Pulling ssDNA-Glycan conjugates through a nanopore with the assistance of a DNA motor

The major challenge of glycan “strand sequencing” is how to stretch the glycan chain and drive it translocate through the nanopore in a velocity-controlled manner, due to lack of driving force. To overcome the limitations, we modified the non-reducing end of the glycan chain by adding charged sialic acid residues and a 30-nucleotide polyT chain (the *lead* strand), thereby increasing the glycan’s negative charge. (Fig 1a, Tab S1) This modification enhanced the driving force of the complex in the electric field, facilitating both the translocation and stretching of the conjugate before and after it passes through the nanopore. Additionally, another ssDNA strand (the *handle* strand, abbreviated as NT89) was conjugated to the glycan chain’s reducing end for binding the TGA motor protein (Hel308 family) to control the translocation speed of the glycan complex (Tab S1) and annealing the *tether* strand with a 3’ cholesterol modification (Tab S1) to adjust the event frequency. Here a poly-LacNAc hepta-saccharide glycan chain-NP-3 (Fig 1a) was used as a model to explore the feasibility of nanopore-based glycan strand sequencing.

**Figure 1.**
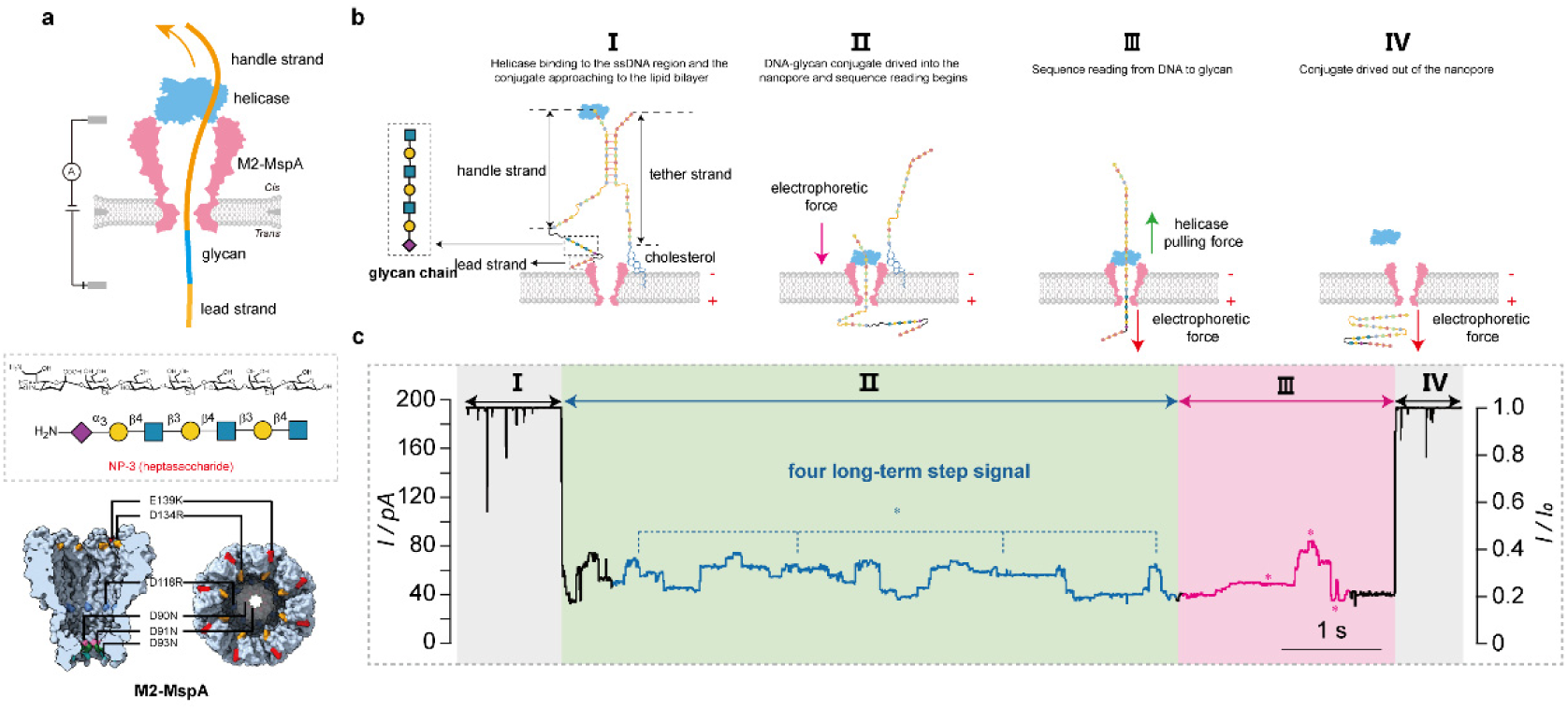
Glycan sequence reading using a nanopore. (a) (upper) Principle of nanopore-based glycan strand sequencing. The model glycan (colored cyan) was conjugated with two ssDNA on both ends, the *lead* strand (polyT, 30nt, colored orange) was used to enhance the electrophoretic force for glycan’s translocation and stretch while the *handle* strand (NT89, 89nt, colored orange) was for the binding of TGA motor (colored blue). The ssDNA-glycan conjugate could be pulled through the M2-MspA nanopore (colored pink) to initiate the sequencing with the control of the TGA motor under +180 mv applied voltage. (middle) Chemical structure and schematic symbol of NP-3. (bottom) Side view and bottom view of M2-MspA with several mutations based on WT-MspA. (b). Schematic view of a single sequencing event. (Ⅰ). helicase bound to the ssDNA region of the *handle* strand and ssDNA-NP-3 conjugate approached the nanopore with the assistance of the *tether* strand modified with cholesterol at the 3’ end (Ⅱ). ssDNA-NP-3 conjugate was driven through the nanopore under +180 mv applied voltage and sequence reading began under the control of TGA motor (Ⅲ). Glycan sequence reading began with the movement of the conjugate. (Ⅳ). TGA motor was dissociated from the conjugate as the linker approached the motor, thus ending the sequencing event. (c) Representative current trace of a single sequencing event. Signals of NT89 were colored in blue and four long-term platform signals were marked with asterisks. Signals related to NP-3 were colored in red and three asterisks refer to the long-term platform signal, the highest current level, and the lowest current level, respectively. M2-MspA nanopore, ssDNA-glycan conjugate, and ATP were all added to the *cis* side. All experiments were performed in symmetric salt buffer (400 mM KCl, 5 mM MgCl_2_, 10 mM HEPEs, pH 8.0) under +180 mv applied from the *trans* side.

By convention, the electrode grounding chamber of the measurement device was defined as the *Cis* chamber, with the opposite chamber defined as the *Tran*s chamber. Positive voltage was applied to the *Trans* side, with M2-MspA (abbreviated as MspA) proteins added to the *Cis* side (Fig.1a). After MspA formed a stable pore, a current around 170 pA could be recorded under +180 mv applied from the *trans* side in a symmetric salt solution (400 mM KCl, 5 mM MgCl_2_, 10 mM HEPEs, pH8.0), after which the *handle* strand-hepta-saccharide-*lead* strand conjugate (abbreviated as NT89-NP-3-polyT)-TGA mixture is added to the *cis* side together with ATP. We found that different from nanopore reads of DNA, ratcheting the conjugate through the nanopore generated a distinct steplike pattern in the ion current. These events were classified into four phases: Stage I, Stage II, Stage III, and Stage IV. (Fig 1b, 1c) In the stageⅠ, before the conjugate-TGA complex occupied the MspA nanopore, the current remained at the open level, with occasional spikes corresponding to the fast movement of ssDNA through the nanopore. No significant changes in current were observed during this phase. When the conjugate was captured by the MspA nanopore and moved under the control of the TGA motor, the current plummeted to a relatively low level, with the I_res_/I_0_ ranging from 20% to 50% approximately, which cannot be observed when TGA&ATP were not added. (Fig 1c, Fig S1) This process consists of two stages (colored in blue and red in Fig 1c) according to the overall current level. The former stage has four long-term platform signals (colored in blue and marked with an asterisk in Fig 1c) with the I_res_/I_0_ lower than 40%, defined as stageⅡ, which we speculated as the translocation of NT89 under the control of TGA motor, as demonstrated by single handle strand signal. (Fig S1) The conjugate and single NT89 shared the same four long-term platform signals. Besides, the platform signal was altered when NT89 was changed to NT89-2, confirming the belongings of this stage. (Fig S2) After this stage, a significant fluctuation in current was observed, characterized by a surge to a long platform (colored in red and marked with an asterisk in Fig 1c) and several stepwise signals followed by the highest level (colored in red and marked with an asterisk in Fig 1c). The current then dropped to the lowest level (colored in red and marked with an asterisk in Fig 1c), before briefly recovering to an I_res_/I_0_ value near that of polyT (I_res_/I_0_ around 20%). This phase, colored in red in Fig 1c, probably corresponds to the translocation of the NP-3 glycan chain. The current exhibited distinct stepwise features and high reproducibility, with I_res_/I_0_ exceeding 40%, which we identified as Stage III. (Fig 1c, Fig S3) The signal patterns observed in the NT89-polyT control conjugate, which lacks NP-3, were similar to those of NT89-NP-3-polyT but did not exhibit the distinct stepwise features associated with NP-3. (Fig S4) This confirmed that Stage III was specifically related to the NP-3 glycan chain. Finally, as the linker part of the conjugate approached the TGA motor, dissociation of the conjugate from the motor occurred, resulting in a return to the open current level, consistent with Stage IV (Fig. 1c). This transition marked the final phase of the event, where the current signal returned to baseline. These observations suggested that the ssDNA-glycan conjugates translocate through MspA nanopores, with sequential current events delineating the controlled passage of glycans facilitated by motor proteins.

### Separation of glycan signatures

To investigate factors that affect NP-3 step-by step signals, we carried out two sets of experiments. First, we conjugated a *handle* strand-glycan complex with different lengths of *lead* strands. As a signal of NT89-NP-3-polyT ended up with a level corresponding to the level of polyT (a flat signal with I_res_/I_0_ near 20% as described previously^24, 32^, a curious question is how many dTMP (deoxythymidine monophosphate) passed through constriction of MspA nanopore when a single sequencing event ended. Accordingly, a series of polyT with different lengths (10nt, 30nt) were designed and used to synthesize NT89-NP-3-polyT. We then acquired, and compared signals from two NT89-NP-3-polyT conjugates and verified that NT89-NP-3-polyT (30nt) and NT89-NP-3-polyT (10nt) share nearly the same signal. (Fig 2a). The only difference is that signals of conjugate with a longer polyT last longer (colored in blue and marked with asterisks in Fig 2a). For MspA, four nucleotides near the constriction determine a step signal^32, 36^ so this means at least four dTMPs stay around the constriction if polyT signals are observed. Since NT89-NP-3-polyT with the length of polyT shortened to 10-mer showed intact NT89, NP-3 signal, and a flat signal polyT signal (I_res_/I_0_ near 20%), 8 dTMPs passed through the constriction at most when a single sequencing event ended. Second, different linkers were constructed to explore the influence of the linker on the glycan step-like signal. The linker near the non-reducing end of NP-3 was changed from SMCC to DBCO (Fig 2b, the corresponding NP-3 were abbreviated as NP-3-SMCC, NP-3-DBCO, respectively, in this section). As described above, a signal of NP-3-SMCC will initially increase to the highest level through a few steps, then decrease to the lowest level via a few steps, and finally step back to the polyT level. However, the lowest level disappeared when SMCC was changed to DBCO. (Fig 2b, Fig S5) We speculated that the conformation of SMCC may lead to a larger occupied volume around the constriction than DBCO do, thus making NP-3-SMCC reveal another lowest level compared with NP-3-DBCO. Besides, the current levels of each step of NP-3-DBCO were lower than that of NP-3-SMCC (for the highest I_res_/I_0_ value, 43% for NP-3-SMCC while 37% for NP-3-DBCO) and this could be explained by the hypothesis that the rigid structure of SMCC may prevent the formation of secondary structure of NP-3 so NP-3-SMCC has an overall higher current level. Nevertheless, both of the conjugates shared the steps before the lowest level, which indicated steps after the lowest level may be mainly determined by the linker and polyT, and steps before the lowest level may be mainly determined by NP-3. Now that we have explored the relationships between the raw step signals and the conjugate sequence, to accurately describe the current signal, we extracted step maps of over 300 events, analyzing the occurrence frequency of all the steps, discarding those with occurrence frequency lower than 50%, and finally identified nine steps from original fourteen steps as the representative step signals for NP-3. (Fig 2c-e) The I_res_/I_0_ value for each step is 19.8%, 24.3%, 31.5%, 38.8%, 43.1%, 40.6%, 33.8%, 17.1%, 19.6%, respectively.

**Figure 2.**
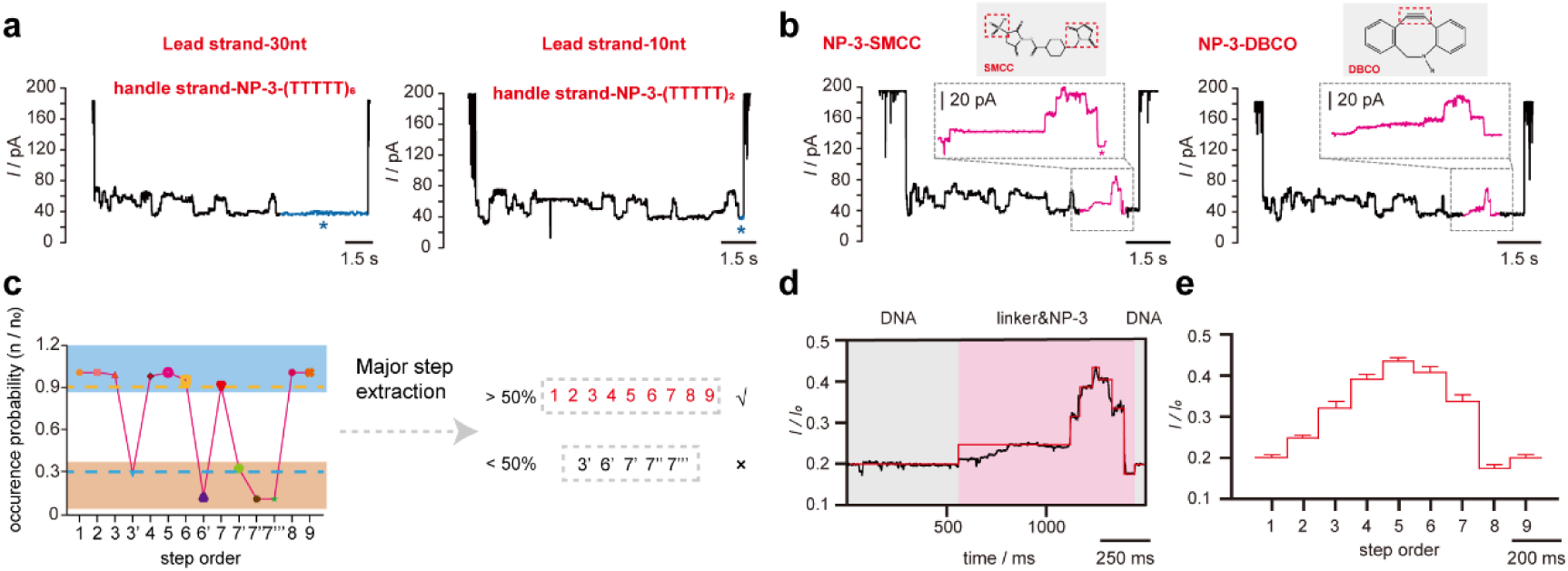
Exploration of conjugate sequence’s influence on the step signature. (a) (left) Representative current trace of NT89-NP-3-polyT (30nt) and (right) NT89-NP-3-polyT (10nt). Signals of polyT were colored in blue and marked with asterisks. (b) (left) Representative current trace of NP-3-SMCC and its glycan signature signal. The lowest current level was marked with an asterisk (n=274). (right) Representative current trace of NP-3-DBCO and its glycan signature signal(n≥15). (c) Frequency distribution scatter plot of steps of NP-3-SMCC (n=354). Steps with frequency over 0.5 were marked with a blue background and extracted as the major step (step 1-9 colored in red) for further analysis while steps with frequency lower than 0.5 were discarded. (d) Current levels of a typical glycan signal of NP-3. Signals of NT89 and polyT at both terminus of the signal were marked with a gray background while the NP-3-related signal was marked with a pink background. (e) Step-by-step signals of NP-3 extracted from multiple sequencing events (n=274). The nine step order corresponds to nine major steps identified in Fig 2c. M2-MspA nanopore, ssDNA-glycan conjugate and ATP were all added to the *cis* side. All experiments were performed in symmetric salt buffer (400 mM KCl, 5 mM MgCl_2_, 10 mM HEPEs, pH 8.0) under +180 mv applied from the *trans* side. The above results further confirmed that the ratcheting motion of the glycan was successfully achieved, generating a characteristic signal pattern corresponding to the sequence of the glycan chain.

### Determination of Glycan Sequencing Read Lengths in Nanopores

To assess the length of glycan sequencing using nanopores, we first characterized the maximum read distance between the helicase and the nanopore constriction. For the M2-MspA nanopore, this distance is approximately 70 Å^37^, which is sufficient to allow the translocation of 7 to 14 monosaccharide units. To experimentally verify this theoretical limit, we synthesized a series of ssDNA-glycan conjugates with varying glycan lengths, specifically NP-3, NP-6, and NP-9, corresponding to seven, nine, and eleven monosaccharide units, respectively. (Fig 2b, Fig3a) These glycans were subjected to nanopore analysis, and their translocation was monitored and reflected by the characteristic polyT-induced current signature, which serves as an indicator of the successful passage of the glycan part through the pore. It is noteworthy that both ssDNA-NP-3 conjugate and ssDNA-NP-6 conjugate ended up with a current level near that of polyT (Fig 1c, Fig 3b, Fig S6, Fig S7) marked with an asterisk), which meant that both NP-3 and NP-6 passed through the constriction completely before a single sequencing event ended. However, when it comes to ssDNA-NP-9 conjugate, the ending current did not decrease back to the polyT level (Fig 3b, Fig S8, Fig S9), suggesting that NP-9 might just partly pass through the constriction when the sequencing event ended. The results revealed that glycan chains shorter than eleven monosaccharide units could be effectively resolved, with distinct stepwise current patterns observed for NP-3 and NP-6. Notably, the observed current blockages were consistent with the theoretical predictions based on the pore’s constriction size, confirming that the read length is limited by the physical dimensions of the nanopore and the translocation process. (data not shown) These results not only validate the maximal read length for glycan of the M2-MspA nanopore system but also provide a reliable method for determining glycan length and structure, advancing the potential of nanopore-based glycan sequencing. Similarly, we identified ten steps from the original thirteen steps as the representative steps for NP-6. (Fig 3c-e) The I_res_/I_0_ value for each step is 19.7%, 23.5%, 21.2%, 35.7%, 39.8%, 41.7%, 37.4%, 32.3%, 16.4%, respectively. The step differences between NP-3 and NP-6 not only lies in the number but also the amplitude and transition characteristics. For NP-3, the step signals increase continuously to step 5-the highest level (I_res_/I_0_ value 43.1%) then decrease to step 8-the lowest level (I_res_/I_0_ value 17.1%), and finally increase back to step 9-the polyT level (I_res_/I_0_ value 19.6%). However, for NP-6, the step signal first increase to step 2-the first peak level (I_res_/I_0_ value 23.5%), then decreases to step 3-the first lowest level (I_res_/I_0_ value 21.2%), then increase to step 6 the highest level (I_res_/I_0_ value 41.7%), finally decrease to step 9-the polyT level (I_res_/I_0_ value 16.4). These results showed that with the assistance of DNA motor protein, discrete step-by-step signals of glycans can be obtained and lengths of different glycans can be read via this system.

**Figure 3.**
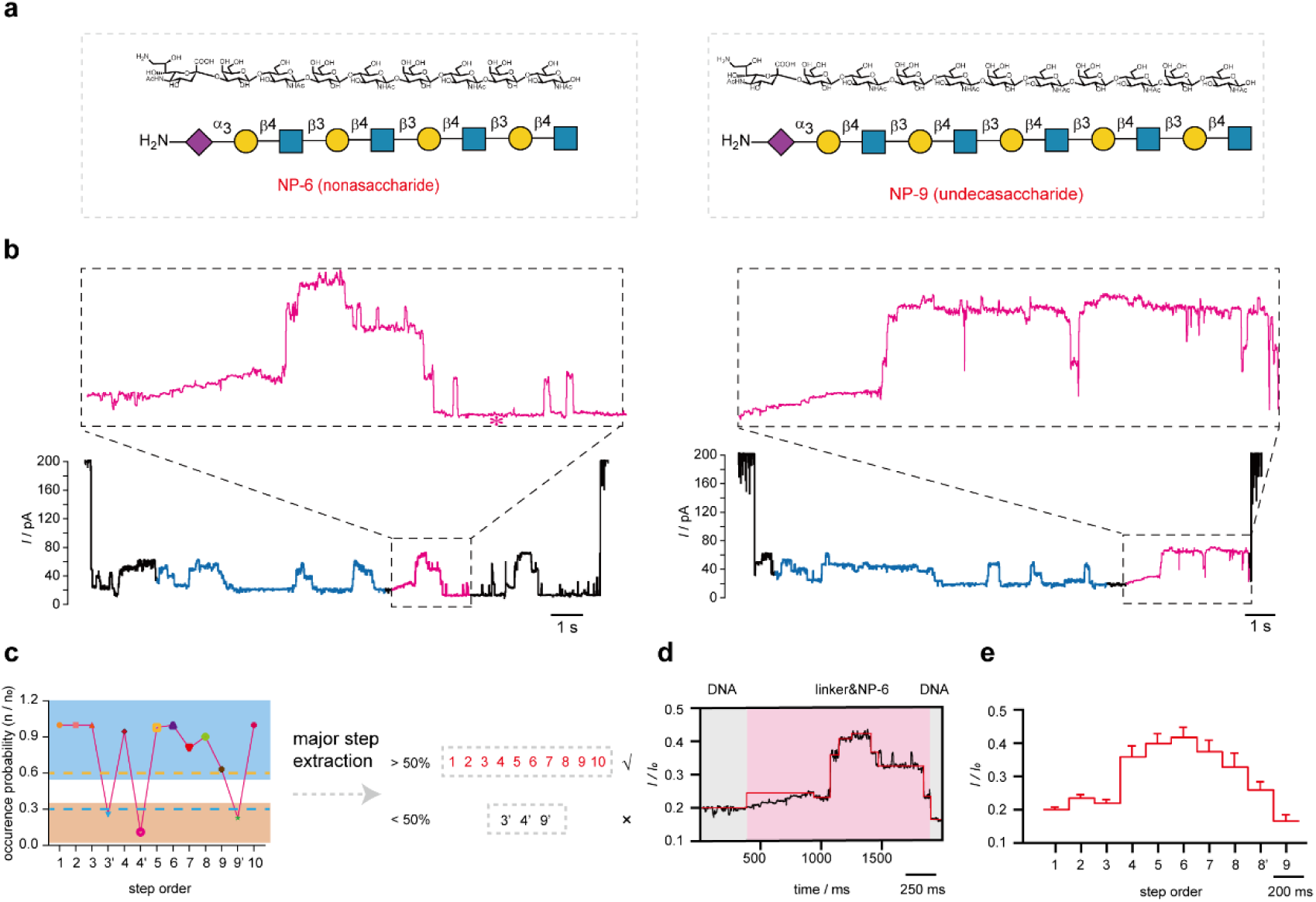
Identification of glycan reading length by M2-MspA nanopore. (a) (left) Chemical structure and schematic symbol of NP-6 and (right) NP-9. (b) (left) Representative current trace of 5T89-NP-6-polyT and the magnifying glycan signal (n=109). The 5T89 and NP-6 signals were colored in blue and red, respectively. The asterisk refers to the current level of polyT. (right) Representative current trace of 6T89-polyT and the magnifying glycan signal (n≥15). The 6T89 and NP-9 signals were colored in blue and red, respectively. (c). Frequency distribution scatter plot of steps of NP-6 (n=298). Steps with a frequency over 0.5 were marked with a blue background and extracted as the major step (step 1-10 colored in red) for further analysis while steps with a frequency lower than 0.5 were discarded. (e) Current levels of a typical glycan signal of NP-6. Signals of 5T89 and polyT at both terminus of the signal were marked with a gray background while the NP-6-related signal was marked with a pink background. (f) Step-by-step signals of NP-6 extracted from multiple sequencing events (n=109). The ten-step order corresponds to ten major steps identified in Fig 3c. M2-MspA nanopore, ssDNA-glycan conjugate and ATP were all added to the *cis* side. All experiments were performed in symmetric salt buffer (400 mM KCl, 5 mM MgCl_2_, 10 mM HEPEs, pH 8.0) under +180 mv applied from the *trans* side.

### Differentiation of Glycan Modification Units in Nanopores

To investigate the influence of glycan modifications on nanopore translocation profiles, we synthesized a series of sulfated glycan conjugates using NP-3 as a template. Specifically, we introduced sulfate groups at a primary hydroxy group of GlcNAc residue at various positions of NP-3, generating three distinct conjugates: NP-23 (sulfated at the 3’ position), NP-18 (sulfated at the 5’ position), and NP-26 (sulfated at both the 3’ and 5’ positions). (Fig 4a-c) Nanopore translocation measurements revealed that, compared to the unmodified NP-3, all sulfated conjugates exhibited a notable reduction in the amplitude of the long-term platform signals. This attenuation suggests that sulfation alters the glycan’s conformational dynamics during translocation, potentially due to steric hindrance or electrostatic interactions with the nanopore. Compared with NP-3, the current level of its first half part corresponding to the long-term platform of NP-23 decreased significantly from 24.3% to 20.1% while the last half part decreased from 24.3% to 22.2%. (Fig 4a, marked with asterisks, Fig S10) This decrease may be explained by the speculation that this level especially the first half part (2*) serves as the time window for sensing the third monosaccharide of the hepta-saccharide glycan chain and the introduction of the sulfonic group, which is more bulky, exacerbates the current blocking effect within the nanopore, thus leading to a significant decrease in the current level. Opposite from 3’ sulfated product NP-23, the last half part of the long-term signal of NP-18 decreased greatly from 24.3% to 16.9% while the first half part decreased from 24.3% to 21.7%. (Fig 4b, marked with asterisks, Fig S11) This phenomenon indicates this level especially the last half part (2**) might serves as the major time window for sensing the 5’ monosaccharide and sulfation of this site inevitably increased the blockage ratio. Notably, NP-26, which was doubly sulfated, displayed counterintuitive changes. Specifically, the decrease in 2* and 2** was from 24.3% to 22.1%, which is higher than that of NP-23 (20.1%), and the decrease in 2** was from 24.3% to 18.0%, which is higher than that of NP-18 (16.9%) (Fig 4c, marked with asterisk, Fig S12). This may be attributed to the enhancement of electrostatic force caused by double-sulfation, which could make the glycan chain more stretched so the current level could be higher. These results suggest that nanopore-based sequencing can be employed to detect and differentiate glycans based on their modification patterns, providing valuable insights into the role of specific modifications in glycan structure and function.

**Figure 4.**
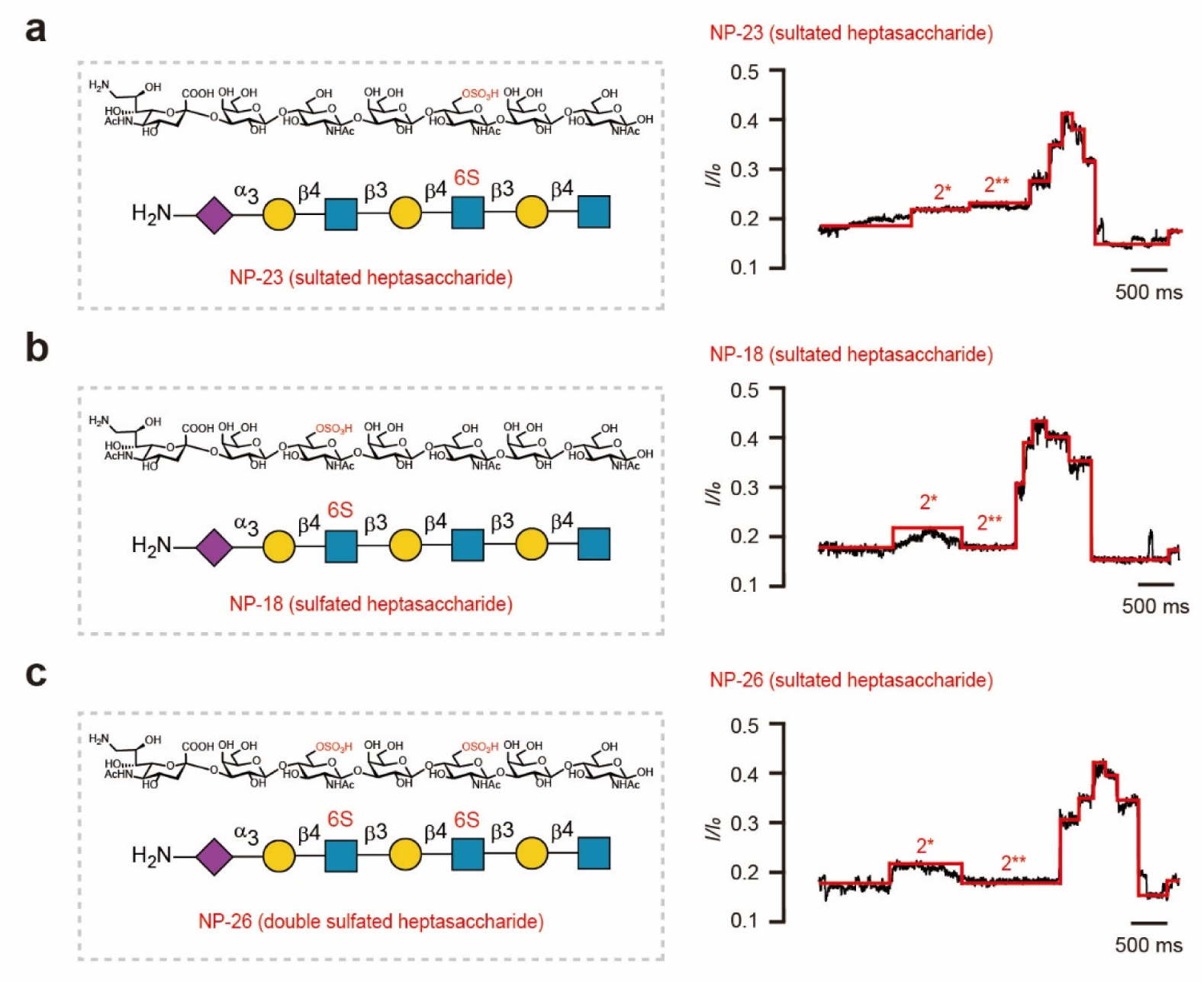
Discrimination of glycans with distinct modification modes. (a) (left) Chemical structure and schematic structure of NP-23. NP-23 was generated by sulfation on the third monosaccharide unit of NP-3. The sulfated site and sulfonic acid group were colored in red. (right) Current levels of a typical glycan signal of NP-23. (n≥15) The black trace was the raw signal and the red line was manually depicted step according to each step level. 2* and 2** were obviously-altered steps after sulfation that corresponded to the long-term platform signal shown in Fig 1c (marked with a pink asterisk). (b) (left) Chemical structure and schematic structure of NP-18. NP-18 was generated by sulfation on the fifth monosaccharide unit of NP-3. The sulfated site and sulfonic acid group were colored in red. (right) Current levels of a typical glycan signal of NP-18. (n≥15) The black trace was the raw signal and the red line was manually depicted step according to each step level. 2* and 2** were obviously-altered steps after sulfation that corresponded to the long-term platform signal shown in Fig 1c (marked with a pink asterisk). (c) (left) Chemical structure and schematic structure of NP-26. NP-26 was generated by sulfation on the third and fifth monosaccharide unit of NP-3. The sulfated site and sulfonic acid group were colored in red. (right) Current levels of a typical glycan signal of NP-26. (n≥15) The black trace was the raw signal and the red line was manually depicted step according to each step level. 2* and 2** were obviously-altered steps after sulfation that corresponded to the long-term platform signal shown in Fig 1c (marked with a pink asterisk). M2-MspA nanopore, ssDNA-glycan conjugate and ATP were all added to the *cis* side. All experiments were performed in symmetric salt buffer (400 mM KCl, 5 mM MgCl_2_, 10 mM HEPEs, pH8.0) under +180 mv applied from the *trans* side.

## Discussion

### Pioneering Strategy for Nanopore-Based Glycan Sequencing Validation

We present here a novel approach to glycan chain sequencing through nanopore technology, utilizing a helicase to control DNA translocation speed, thereby indirectly modulating glycan translocation rates. This strategy achieves unprecedented single-molecule sensitivity, stretching and controlling the movement of glycan chains, and enables glycan signal reading in a linear configuration for the first time. Our method exhibits the ability to resolve individual monosaccharides with high sensitivity and to repeatedly sequence sugar units, modifications, and linkages (Data not shown) within specific sequence contexts. This advancement introduces a viable approach for real-time monitoring of glycan modifications and represents a new paradigm for glycan sequencing technology.

### Development of a Method for Glycan Signal Acquisition and Analysis

To further understand glycan signals, we systematically explored how variables such as linker composition, glycan lengths, and modifications affect the stepwise pattern of the recorded signals. By altering these components, we elucidated preliminary correlations between glycan structure and signal patterns, forming the basis for a more detailed exploration of glycan-DNA-linker signal relationships. Subsequent studies using polyT point mutations or complete substitutions, such as polyT-to-polyA, will enable a more granular analysis. This work lays foundational insights into glycan signal formation mechanisms and supports further development in high-precision glycan sequence analysis.

### Algorithmic Framework for Stepwise Recognition of Glycan Nanopore Electrical Signals

Additionally, we established an initial algorithm for recognizing stepwise glycan signals, offering a prototype for accurate glycan sequence mapping. Though still in its developmental stages, this algorithm underpins the establishment of a standardized glycan signal database and opens the door to glycan sequence identification with enhanced specificity and sensitivity. Once optimized, this approach could support high-resolution, single-molecule glycan sequencing.

## Conclusion

Our study achieved a breakthrough in single-molecule glycan sequence reading, demonstrating the first chain-wise reading of glycan signals through controlled translocation through the nanopore. We uncovered key aspects of glycan signal patterning and developed an initial algorithm for glycan sequence recognition. This work advances glycan sequencing by establishing a new method capable of discerning individual sugar units, modifications, and linkage types within their unique sequence contexts. Ultimately, this method may enable the precise fingerprinting and full-length identification of complex glycan molecules. The compatibility of our approach with commercial equipment, combined with its cost-efficiency and adaptability, holds significant promise for single-molecule glycan sequencing and glycomics research. This method could enable highly sensitive detection of glycan structures within proteins, broadening its application scope across fundamental biology, drug discovery, and clinical settings. Our strategy represents an initial step toward achieving real-time glycan modification monitoring and glycan analysis at single-molecule-level.

### Limitations and Future Challenges

Despite these advances, challenges remain. Directly decoding glycan sequences from nanopore signals is complex and requires extensive datasets for glycan sequence training and recognition. Additionally, the structural complexity of glycans poses difficulties in signal resolution, necessitating further optimization of nanopore read-out accuracy. Given that glycan monomer diversity far exceeds that of nucleotides or amino acids, creating a complete mapping of ionic current signatures to individual monosaccharides demands substantial data, which may limit practical implementation. Enhanced signal capture could be achieved by engineering the nanopore or by incorporating charged tags at glycan termini to improve the translocation of neutral sugars. We will subsequently continue exploring the relationships between the glycan sequence and the step signal by extracting glycan signals from diverse structures including distinct linkages and constructing an accurate raw current signal-step signal-glycan sequence map.

## Methods

### Purification of M2-MspA nanopore and TGA protein

First, the M2-MspA plasmid was introduced into Acella receptor cells, which were subsequently uniformly coated on LB solid medium containing kanamycin and inverted and cultured overnight at 37°C to promote the transformation of the plasmid and the growth of the clone. Next, single colonies were selected from the above medium, inoculated into LB liquid medium, and kanamycin was added for selective cultivation to ensure that all cultured bacteria contained the M2-MspA plasmid. After overnight incubation, the bacterial solution was expanded into LB medium, and kanamycin and a small amount of previously cultured bacterial solution were continued to be added, and the culture was incubated until the OD600 value reached 0.6-0.8, and then IPTG was added to induce the expression of MspA M2 protein. Subsequently, the bacteria were collected by centrifugation and resuspended using AEX-A buffer at a mass volume ratio of 1:9 and TritonX-100 was added to a final concentration of 2%. Next, high-pressure fragmentation was performed at less than 6°C to release intracellular M2-MspA proteins, followed by centrifugation and filtration to remove insoluble impurities. Next, the supernatant was initially purified using an anion-exchange column, and the flow-through solution was collected until the UV value decreased to 335 mAU. After that, precipitation was carried out by adding saturated ammonium sulfate, and the supernatant was removed by centrifugation after settling to obtain the crude extract of M2-MspA protein. The crude extract was re-dissolved in a 1:3 mass-volume ratio and incubated overnight at 4°C to further stabilize the protein structure. On the following day, insoluble impurities and possible denatured proteins were removed by centrifugation and heat treatment, and then the supernatant was concentrated to half volume using an ultrafiltration tube. Finally, the concentrated supernatant was passed through molecular sieve SD200 for final purification to obtain highly pure M2-MspA protein. After determining the protein concentration, glycerol was added proportionally for preservation and placed at -20℃ for subsequent use. The whole preparation process requires strict control of temperature, pH, and ionic strength to ensure the stability and purity of M2-MspA protein.

TGA protein was purified as described previously^24^. In brief, the gene of TGA protein was cloned to the pET15b vector with seven histidine residues on the N-terminus. After transforming the competent cell with TGA plasmid, a single colony on the LB agar medium was picked for sequencing for sequence verification. Then the verified colony was amplified in LB medium from low volume to high volume. The cell precipitation was collected for lysing with a high-pressure homogenizer after induction and amplification and finally purified with Ni-NTA, Heparin column, and SD200 size exclusion chromatography. The purified protein was verified by SDS-PAGE.

### Conjugation of ssDNA and glycans

For conjugation when the linker near the non-reducing end of the glycan is DBCO, the whole synthesis consists of three steps. First, the handle strand with 5’ DBCO modification is mixed with the glycan, reacting for 2h, after which product 1 (handle strand-glycan) was separated by ethanol precipitation and freeze-drying. Second, product 1 was mixed with benzene-sulfonyl azide, reacting for 2 h, and product 2 (handle strand-glycan conjugate with an azido group at the glycan terminus) was separated in the same way. Finally, product 2 was mixed with a lead strand with 3’ DBCO modification, reacting for 2 h. The product was separated by ethanol precipitation, and free-drying, after which the raw product was purified by HPLC to acquire a pure target product (handle strand-glycan-lead strand). For conjugation when the linker near the non-reducing end of the glycan is SMCC, the first step is almost the same as the above synthesis. After that, handle strand-glycan was mixed with SMCC, reacting for 1h, followed by the desalting process to acquire handle strand-glycan conjugate with a maleimide group at the glycan terminus. Finally, the modified handle strand-glycan conjugate was mixed with the lead strand with 3’-SH modification, reacting for 10 h. The product was purified by 15% Urea-PAGE to acquire pure target product (handle strand-glycan-lead strand). Before analysis, the ssDNA-glycan conjugate is mixed with the tether strand and annealed in a thermocycler with the following program-86℃ 1min, 76℃ 1min, 66℃ 1min, 56℃ 1min, 46℃ 1min, 36℃ 1min, 26℃ 1min, 16℃ 1min. Then the mixture was mixed with a TGA motor for the following experiment.

### Electrophysiology recordings and data analysis

In general, 1,2-diphytanoyl-sn-glycerol-3-phosphocholine (DPhPC) was added to the salt buffer in the chamber separated by a polytetrafluoroethylene (PTFE) membrane with a 30-70 μm pole on the center into two sides, the grounded side (the cis side) and the other side (the trans side), respectively. By adjusting the solution level to a height near the pole with a pipette, a lipid bilayer was formed quickly. Unless otherwise stated, each side contains 250 μl symmetric salt solution (400 mM KCl, 5 mM MgCl_2_, 10 mM HEPEs, pH8.0) with M2-MspA nanopore and analytes added to the cis side under +180 mv applied from the trans side. Once an M2-MspA nanopore was inserted into the lipid bilayer stably, a current of around 170 pA occurred, after which the analytes and ATP were added to the cis, and the corresponding real-time current signal was continuously recorded using Smartnano, sampled at 50 kHz, and filtered at 500 Hz. Then the raw current signal was first filtered with an 8-pole Bessel filter at 100 Hz before analysis. Afterward, the current signal was analyzed by Pynanolab or pClamp 10.2 to acquire step features of each ssDNA-glycan conjugate. The figures were plotted using Graphd Prism 9.

## Acknowledgments

We are grateful to Shanghai Municipal Science and Technology Major Project, the National Science Fund of Distinguished Young Scholars (Grant 81825021), Fund of Youth Innovation Promotion Association (Grants 2019285, 2021333, and 2022077), the National Natural Science Foundation of China (Grants 92169202, 82341058), Fund of Shanghai Science and Technology Innovation Action Plan (Grant 20ZR1474200), Shanghai Rising-Star Program (Grant 22QA1411000), Natural Science Foundation of Shanghai (Grant 22ZR1474000).

## Author Contributions

Z.G., J.B., L.W., and B.X. conceived the project. B.X., Z.G., and J.B. designed the experiments; W.K., P.W., Y.Y. and Y.W. carried out the recordings; X.L.,S.M., Z.Y. and Z.Q. Performed Chemical synthesis. All authors analyzed and discussed the data. B.X., Z.G., J.B. and L.W wrote the manuscript. All authors read and approved the manuscript.

## Notes

The authors declare no competing financial interest.

## ACKNOWLEDGMENT

We are grateful to Shanghai Municipal Science and Technology Major Project, Fund of Youth Innovation Promotion Association (Grants 2019285, 2021333, and 2022077), the National Natural Science Foundation of China (Grants 92169202, 82341058), Fund of Shanghai Science and Technology Innovation Action Plan (Grant 20ZR1474200), Shanghai Rising-Star Program (Grant 22QA1411000), Natural Science Foundation of Shanghai (Grant 22ZR1474000) for financial support.

## Supporting information

**Fig S1.**
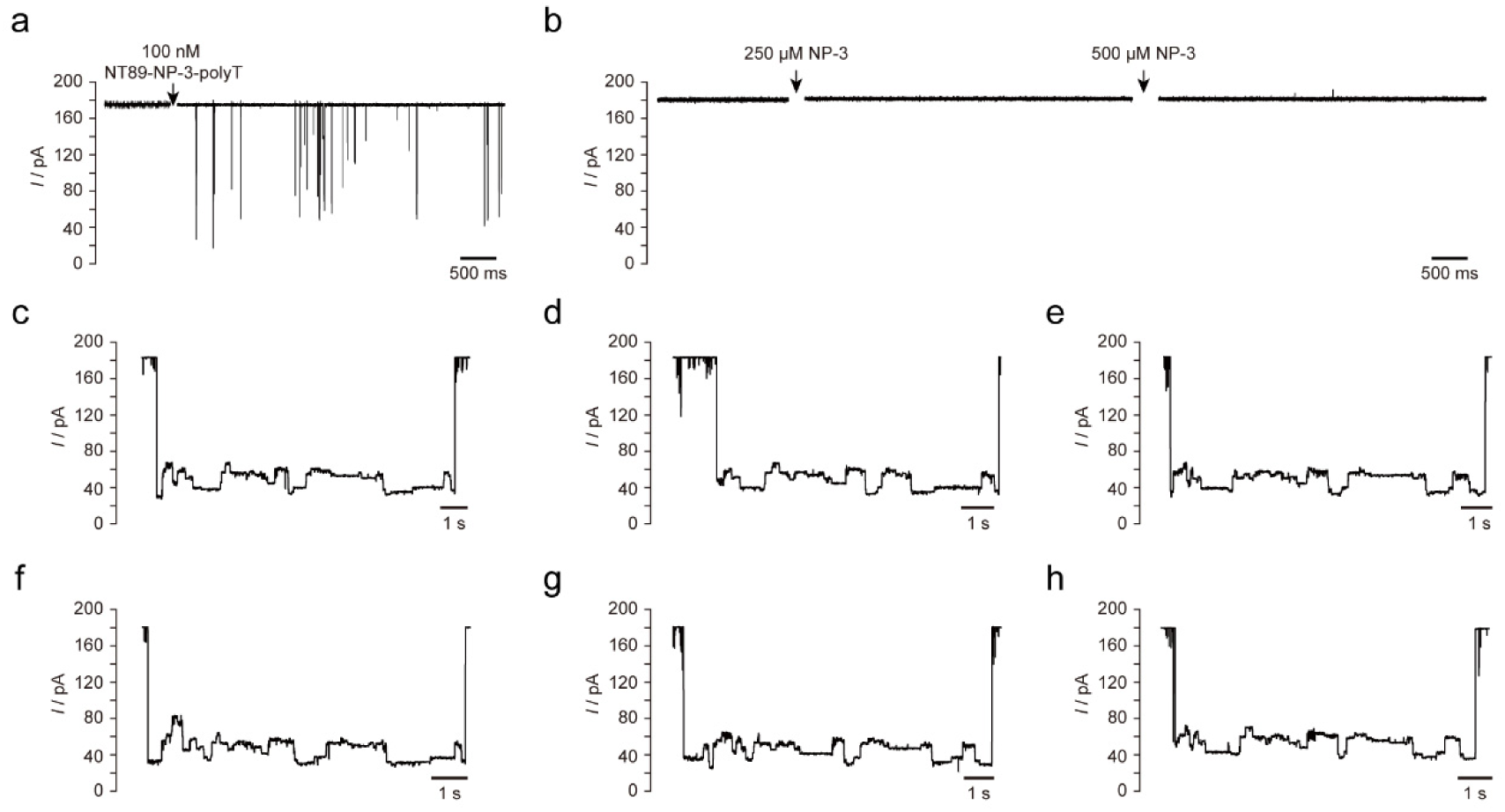
Representative current signal of NT89-NP3-polyT, NP-3, NT89. (a) Representative current trace of NT89-NP-3-polyT without adding TGA motor and ATP. No step-like signals were observed. (n≥3) (b) Representative current trace of NP-3 glycan. No signal was observed even when the concentration increased to 500 μM. (c-h) Reproducibility of NT89 current signal. The analytes were added to the *cis* side. All experiments were performed in symmetric salt buffer (400 mM KCl, 5 mM MgCl_2_, 10 mM HEPEs, pH8.0) under +180 mv applied from the *trans* side.

**Fig S2.**
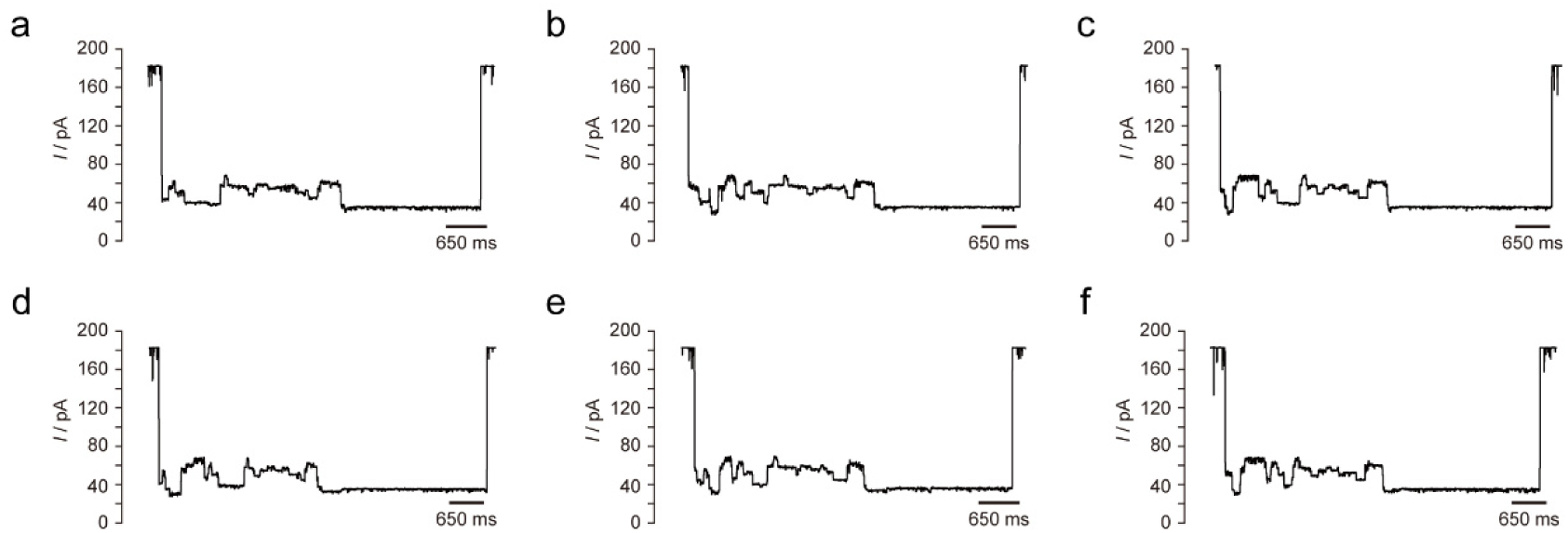
Representative current signals of NT89-2. (a-f) Reproducibility of NT89-2 current signal. The analytes were added to the *cis* side. All experiments were performed in symmetric salt buffer (400 mM KCl, 5 mM MgCl_2_, 10 mM HEPEs, pH8.0) under +180 mv applied from the *trans* side.

**Fig S3.**
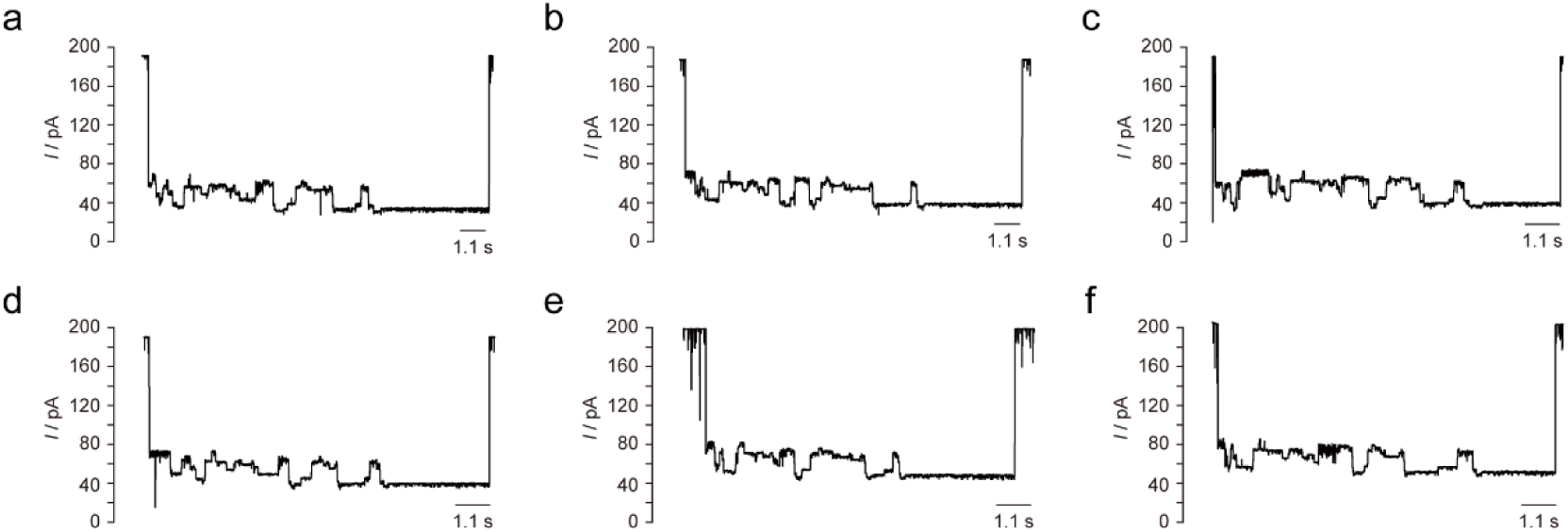
Representative current signals of NT89-polyT. (a-f) Reproducibility of NT89-polyT current signal. The analytes were added to the *cis* side. All experiments were performed in symmetric salt buffer (400 mM KCl, 5 mM MgCl_2_, 10 mM HEPEs, pH8.0) under +180 mv applied from the *trans* side.

**Fig S4.**
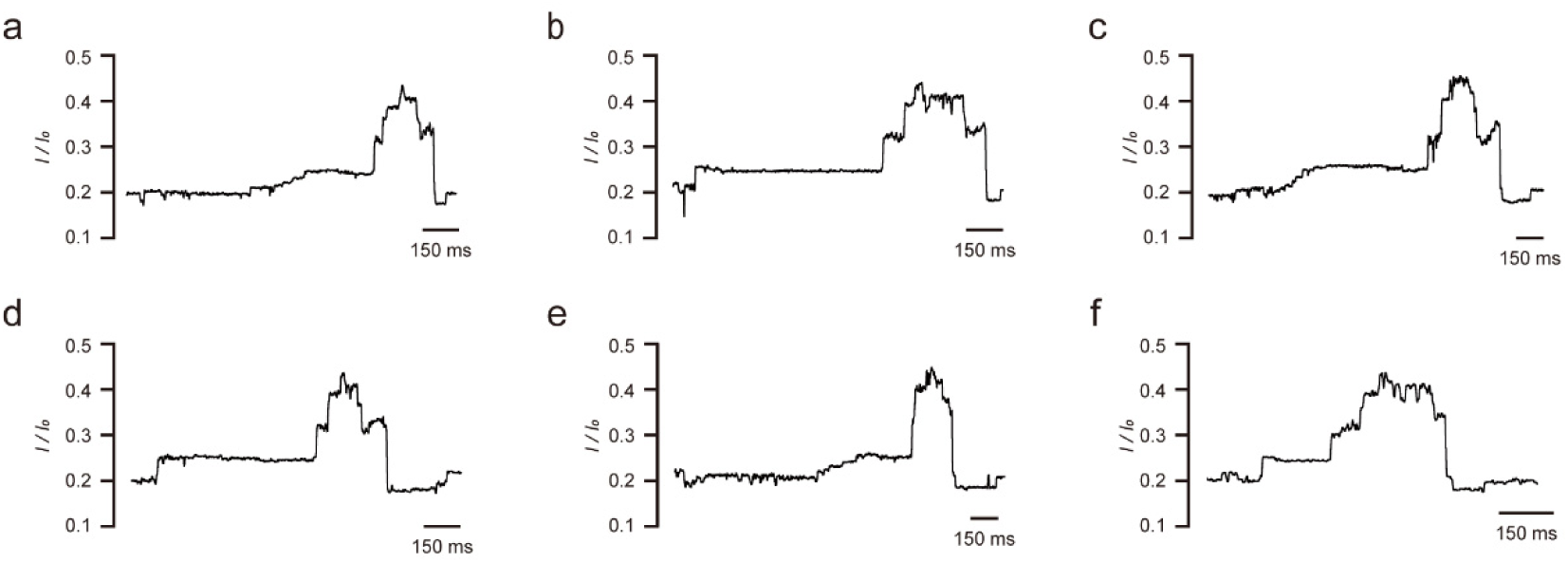
Representative current signals of NP-3-SMCC. (a-f) Reproducibility of NP3-SMCC current signal. The analytes were added to the *cis* side. All experiments were performed in symmetric salt buffer (400 mM KCl, 5 mM MgCl_2_, 10 mM HEPEs, pH8.0) under +180 mv applied from the *trans* side.

**Fig S5.**
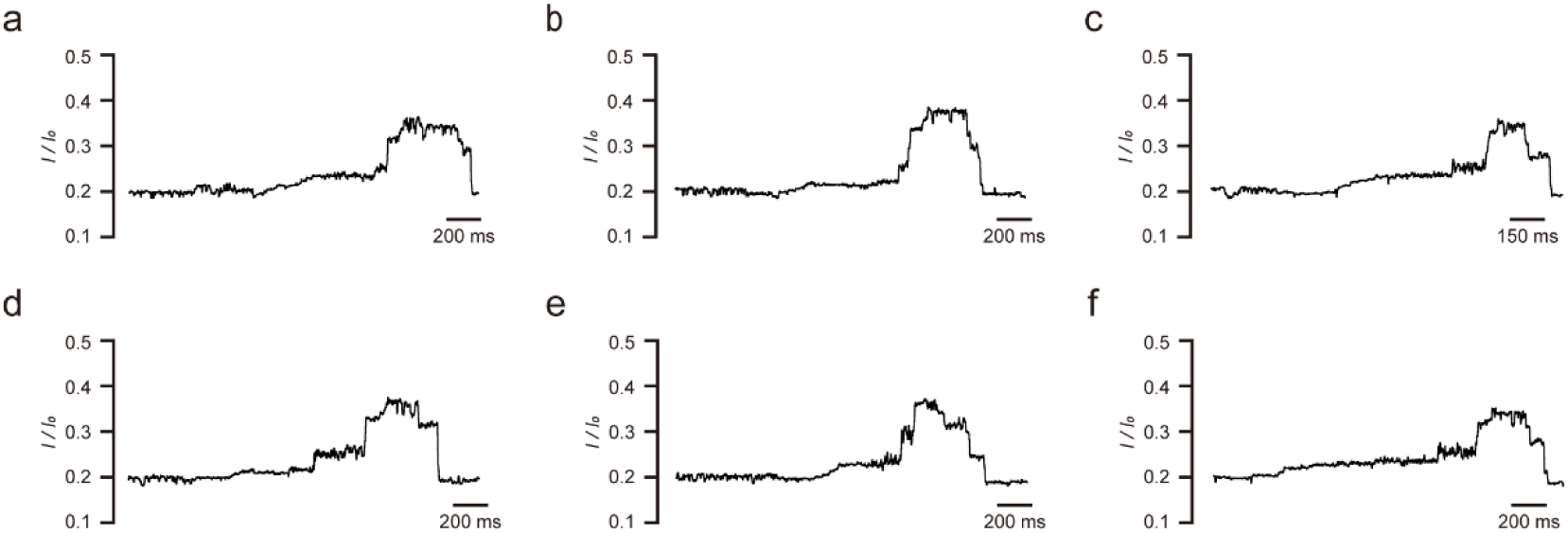
Representative current signals of NP-3-DBCO. (a-f) Reproducibility of NP3-DBCO current signal. The analytes were added to the *cis* side. All experiments were performed in symmetric salt buffer (400 mM KCl, 5 mM MgCl_2_, 10 mM HEPEs, pH8.0) under +180 mv applied from the *trans* side.

**Fig S6.**
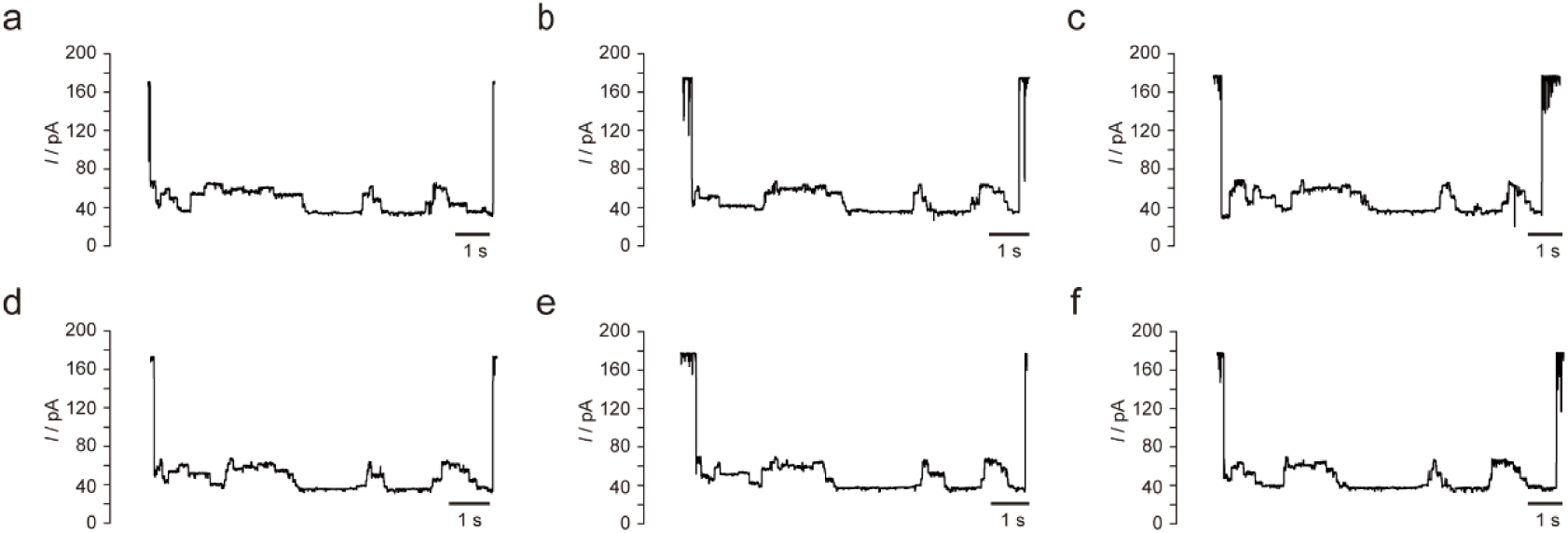
Representative current signals of NT89-3. (a-f) Reproducibility of NT89-3 current signal. The analytes were added to the *cis* side. All experiments were performed in symmetric salt buffer (400 mM KCl, 5 mM MgCl_2_, 10 mM HEPEs, pH8.0) under +180 mv applied from the *trans* side.

**Fig S7.**
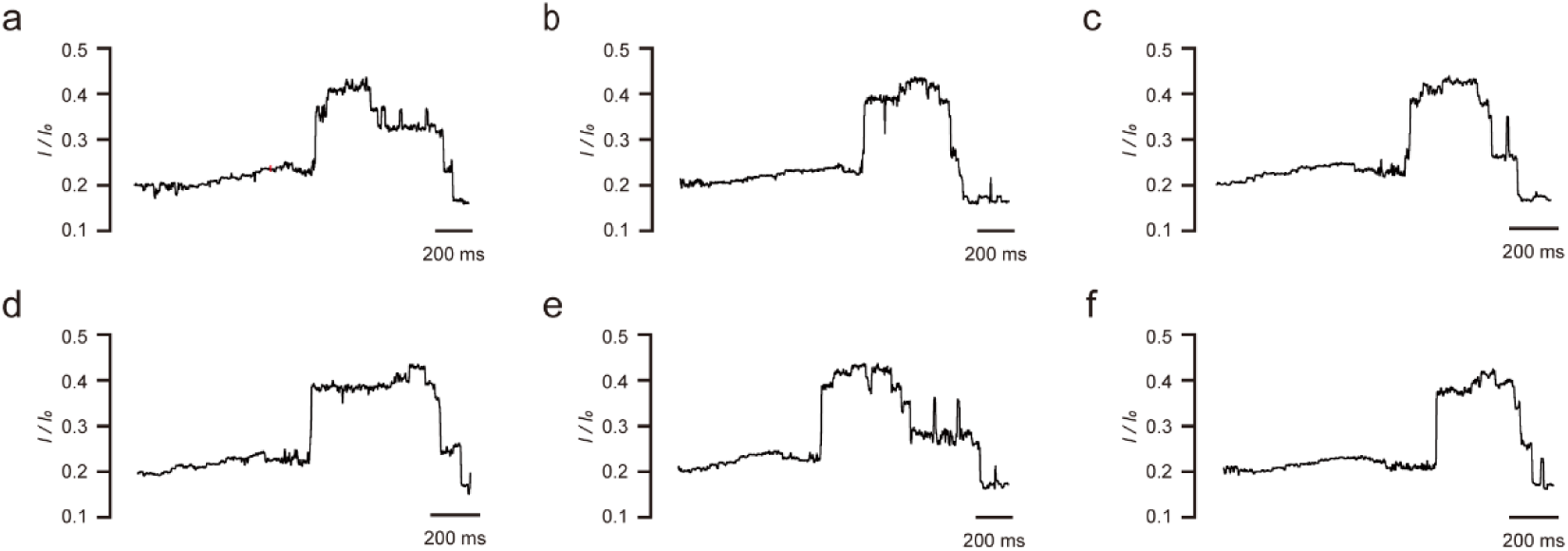
Representative current signals of NP-6-SMCC. (a-f) Reproducibility of NP6-SMCC current signal. The analytes were added to the *cis* side. All experiments were performed in symmetric salt buffer (400 mM KCl, 5 mM MgCl_2_, 10 mM HEPEs, pH8.0) under +180 mv applied from the *trans* side.

**Fig S8.**
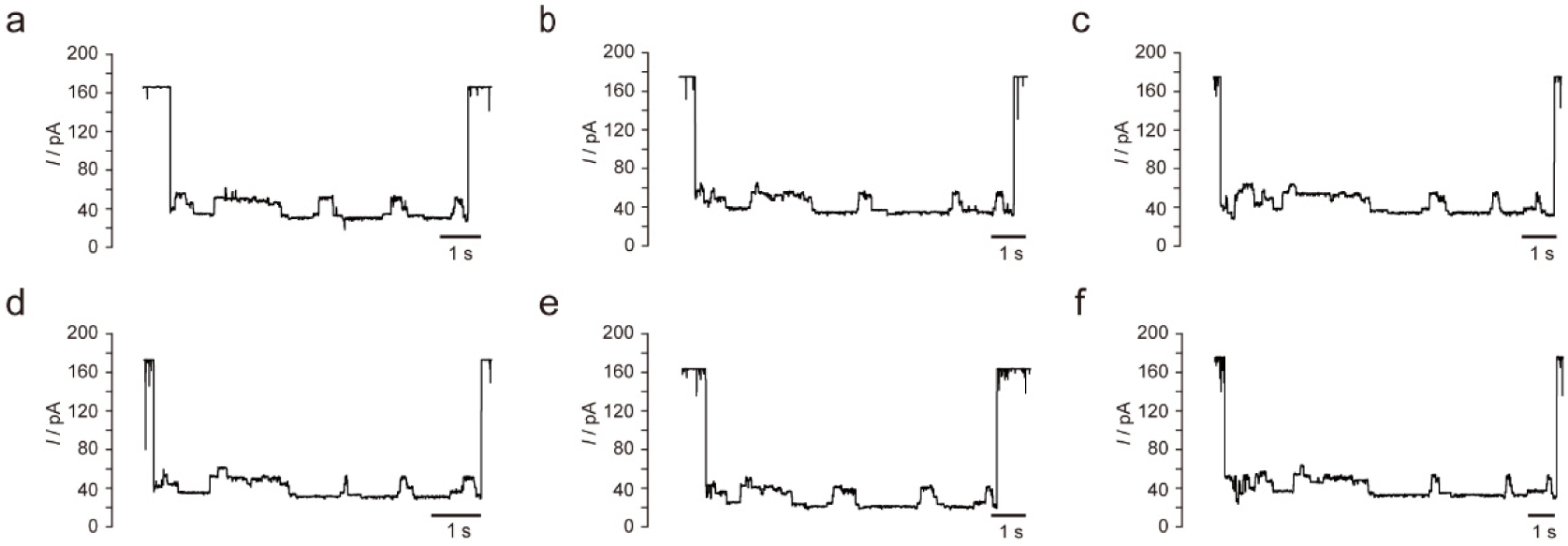
Representative current signals of NT89-4. (a-f) Reproducibility of NT89-4 current signal. The analytes were added to the *cis* side. All experiments were performed in symmetric salt buffer (400 mM KCl, 5 mM MgCl_2_, 10 mM HEPEs, pH8.0) under +180 mv applied from the *trans* side.

**Fig S9.**
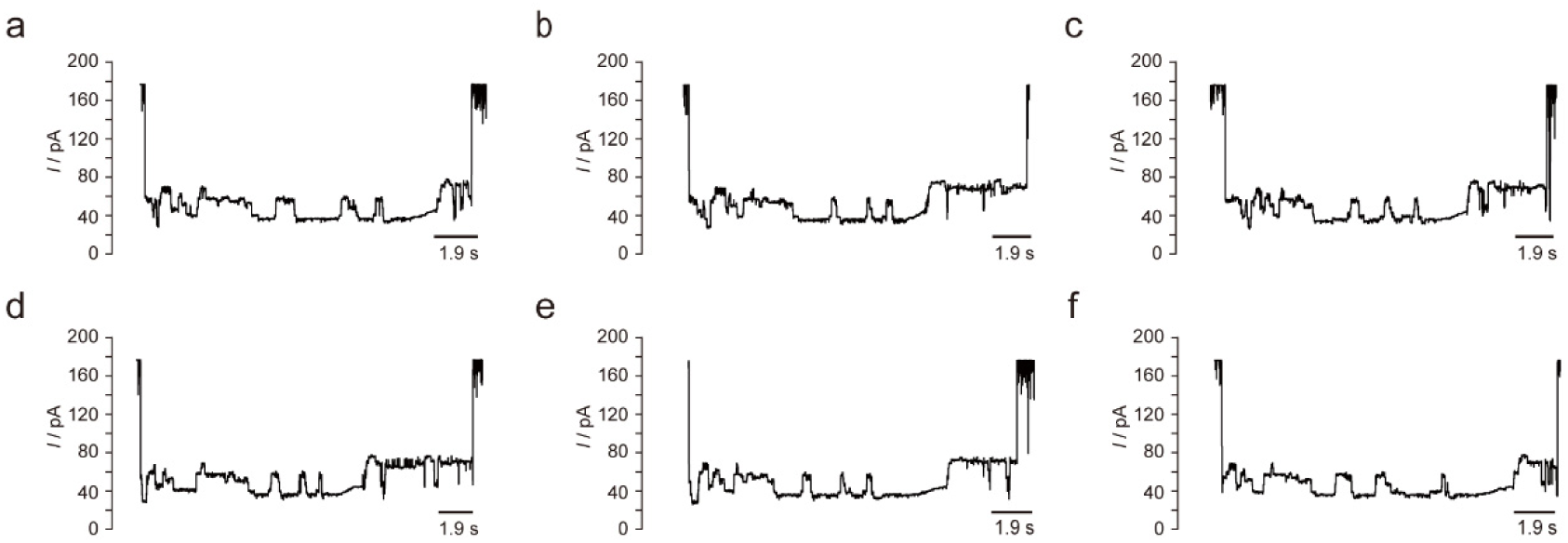
Representative current signals of NP-9-SMCC. (a-f) Reproducibility of NP-9-SMCC current signal. The analytes were added to the *cis* side. All experiments were performed in symmetric salt buffer (400 mM KCl, 5 mM MgCl_2_, 10 mM HEPEs, pH8.0) under +180 mv applied from the *trans* side.

**Fig S10.**
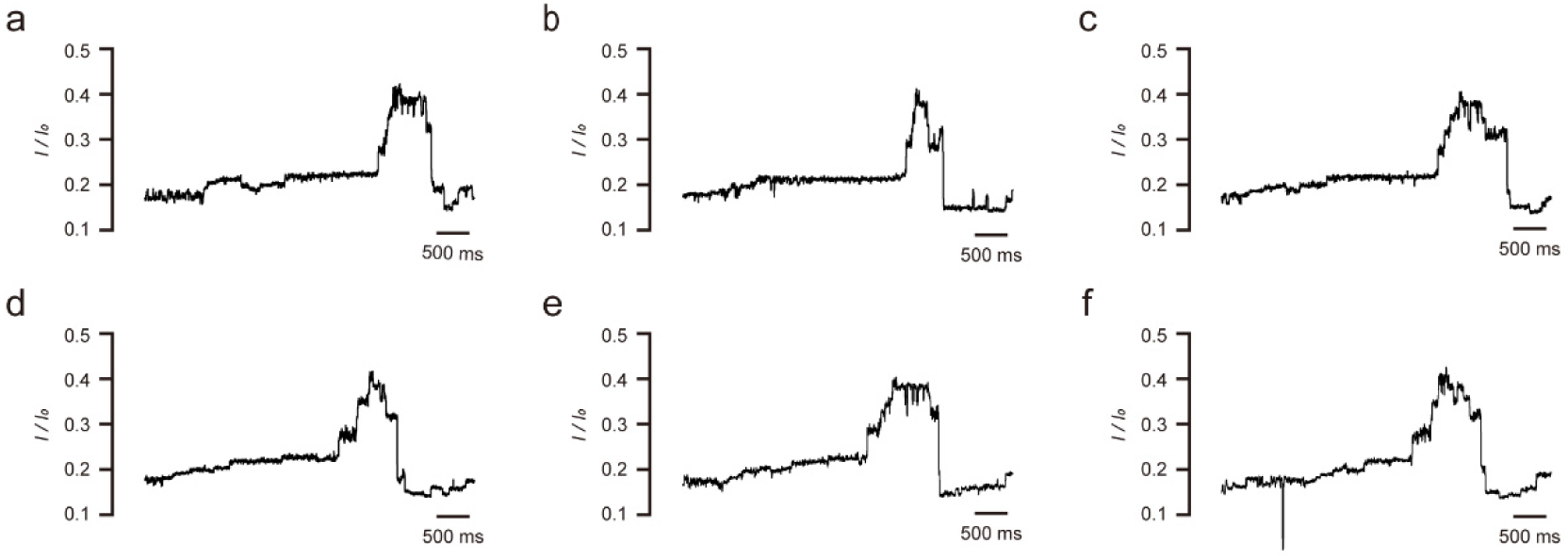
Representative current signals of NP-23. (a-f) Reproducibility of NP-23 current signal. The analytes were added to the *cis* side. All experiments were performed in symmetric salt buffer (400 mM KCl, 5 mM MgCl_2_, 10 mM HEPEs, pH8.0) under +180 mv applied from the *trans* side.

**Fig S11.**
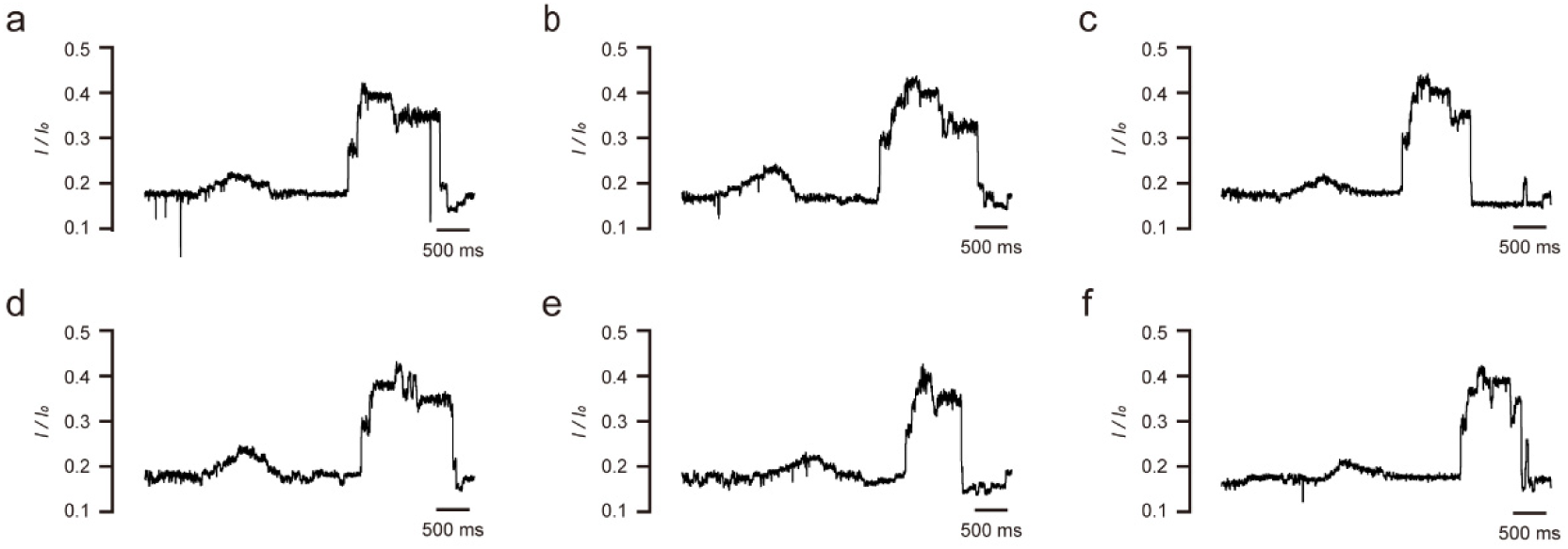
Representative current signals of NP-18. (a-f) Reproducibility of NP-18 current signal. The analytes were added to the *cis* side. All experiments were performed in symmetric salt buffer (400 mM KCl, 5 mM MgCl_2_, 10 mM HEPEs, pH8.0) under +180 mv applied from the *trans* side.

**Fig S12.**
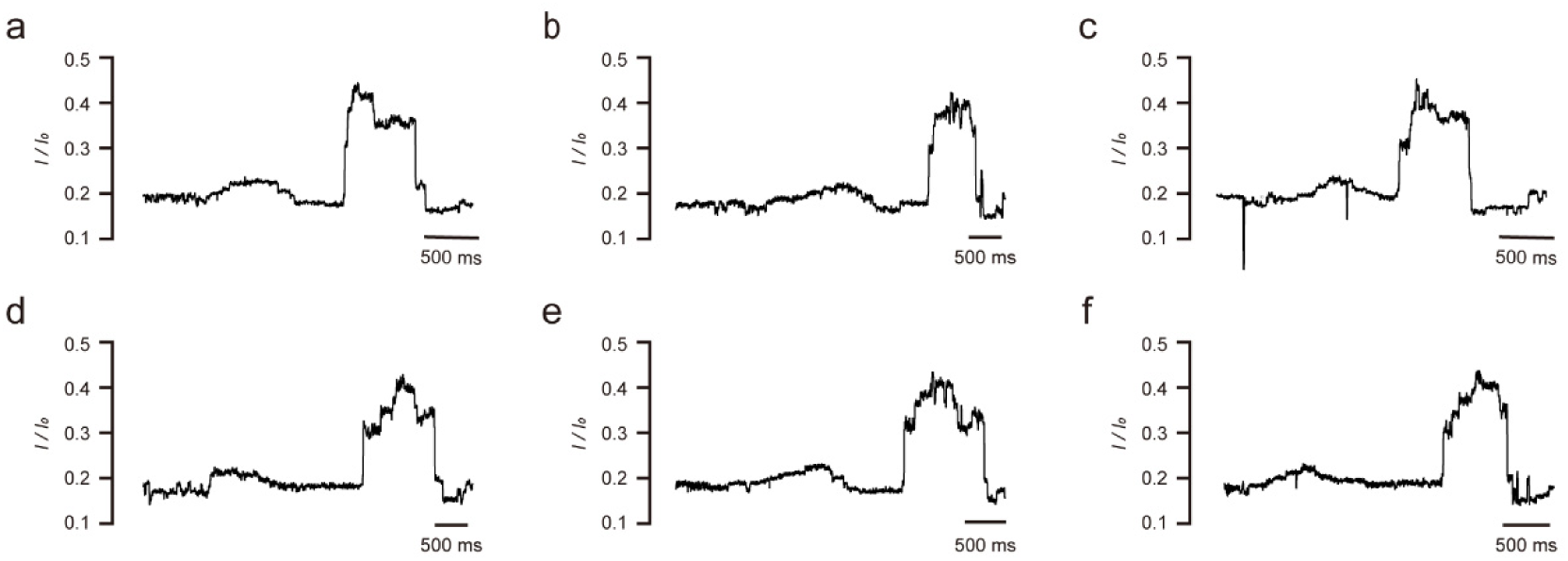
Representative current signals of NP-26. (a-f) Reproducibility of NP-26 current signal. The analytes were added to the *cis* side. All experiments were performed in symmetric salt buffer (400 mM KCl, 5 mM MgCl_2_, 10 mM HEPEs, pH8.0) under +180 mv applied from the *trans* side.

**Table S1.**
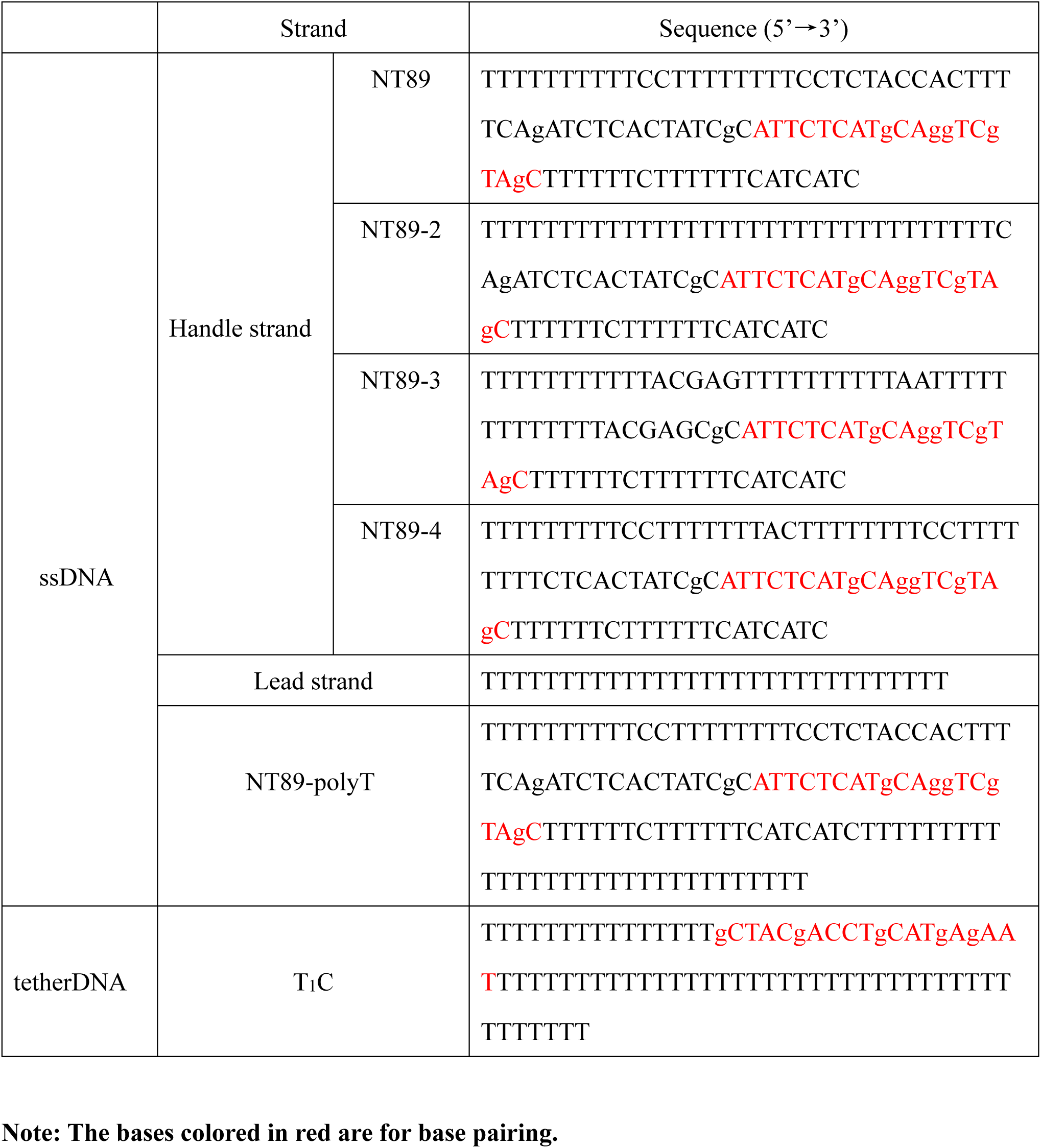
Sequences of single-strand DNA

